# Development of next-generation sequencing-based sterility test

**DOI:** 10.1101/2020.12.27.424516

**Authors:** Jianhui Yue, Chao Chen, Xiaohuan Jing, Qiwang Ma, Bo Li, Xi Zhang

## Abstract

The sterility testing methods described in pharmacopoeias require an incubation period of 14 days to obtain analysis results. An alternative method that can significantly shorten the detection time and improve the accuracy is in urgent need to meet the sterility testing requirements of regenerative medicine products with a short shelf life. In this study, we developed the next-generation sequencing-based sterility test (NGSST) based on sequencing and multiple displacement amplification. The NGSST can be finished within 48 hours with five steps including whole genome amplification, sequencing, alignment, sterility testing report, and microorganism identification. We use RPKM ratio to minorize the influence of environmental bacteria and determine its cutoff based AUC curve. The NGSST showed high sensitivity in reporting contaminates at 0.1 CFU in supernatant of biological product or 1 CFU in cell suspension. Furthermore, we identified microorganisms in 5 primary umbilical cord mesenchymal stem cell samples that were tested positive by BacT/ALERTR 3D. Overall, the NGSST can serve as a promising alternative for sterility testing of biological products.

## Introduction

The current ‘gold standard’ compendial test for pharmaceutical sterility testing first appeared in the British Pharmacopeia in 1932 and has remained fundamentally unchanged ever since. This test requires that product samples be placed in aerobic and anaerobic culture medium for 14 days, during which no growth of microorganisms (bacteria and fungi) should be observed. Although the method can be applied to biopharmaceuticals, it is not perfect fit for regenerative medicine products used in tissue engineering, cell and gene therapies, etc. These special biological products are composed of living cells with relatively small product volume and short shelf life (a few hours to several days), which is challenging to meet the requirements of large sample volume and long-term incubation for sterility testing methods in pharmacopoeias. In particular, the test period of up to 14 days pose a major hurdle for the release of short-lived cellular products. In addition, the pharmacopoeia methods are not effective for the detection of many environmental microorganisms that cannot be cultivated in the culture medium. Therefore, to obtain more comprehensive analysis and shorter test duration, an alternative sterility test method is needed to improve product quality and reduce time and cost for product manufacturing and releasing.

In 2012, FDA revised 21 CFR 610.12 to encourage the use of the more appropriate and advanced testing methods to ensure the safety of biological products [1]. The purpose of the revised requirements of sterility testing is to promote the improvement and innovation of sterility test methods to meet the challenges of new products such as viral and gene therapy (VGT) and cell-based products that may be introduced into the market. Several rapid microbial contamination detection methods have been developed so far, such as ATP bioluminescence, flow cytometry, nucleic acid amplification, solid phase cytometry and so on. However, these methods have certain limitations in sensitivity or accuracy [2–4].

Next-generation sequencing (NGS), also known as high-throughput sequencing, can simultaneously sequence thousands to billions of DNA fragments independently. Recently, the development of metagenomic NGS (mNGS) technology and rapid bioinformatics pipelines allows unbiased detection of pathogens in samples such as urine, cerebrospinal fluid, blood and so on [5–9], which may contain mixed populations of microorganisms. Compared with traditional pharmacopoeia methods, NGS technology have certain advantages, including (1) shorter detection period; (2) unbiased detection and identification of microorganisms; (3) higher sensitivity; (4) no need of cell culturing. However, there are still several challenges in applying NGS technology in sterility testing, such as the lack of systematic validation and bioinformatics pipeline for rapid data analysis that can reduce the bioinformatics processing time from days to several hours [10–11].

Multiple displacement amplification (MDA) is a method of whole genome amplification using very small amounts (<10 pg) of DNA [12]. Currently, most MDA based single cell whole genome amplification kits were used in human or animal cells. There is a lack of research on the performance of using MDA for whole genome amplification of microbes such as bacteria and fungi.

In this study, we applied the mNGS/MDA in sterility testing and developed the next-generation sequencing-based sterility test (NGSST). The NGSST has five steps including whole genome amplification, sequencing, alignment, sterility test report, and microorganism identification. To minorize the influence of environmental bacteria, we used RPKM ratio with cutoff 2.45 to determine microorganism contaminates. The NGSST showed high sensitivity in reporting contaminates at 0.1CFU in supernatant of biological product or 1 CPU in cell suspension. Some of these 0.1 CFU microorganism sample were directly or indirectly (cultured in medium for 14 days) validated by BacT/ALERTR 3D. The NGSST was applied and identified microorganism in 5 primary umbilical cord mesenchymal stem cell samples that were tested positive using traditional pharmacopoeia method.

## Materials and Methods

### Strains

Several strains of microorganisms representing aerobic, anaerobic, Gram-positive, Gram-negative, yeast and fungi were used in this study to assess the novel method of sterility testing based on NGS. These microorganisms grow slowly, need a large amount of nutrition, and have a low content of gas chromatography. They were purchased from national center for medical culture collections (CMCC) in China, typical culture preservation centers in the United States (ATCC) and Japan Collection of Microorganisms (JCM) (Table S1).

To determine the cell viability, Staphylococcus aureus, Pseudomonas aeruginosa, Bacillus subtilis and Escherichia coli were cultured in tryptic soy broth (TSB) at 35°C for 3-5 days, while Clostridium sporogenes and Fusobacterium nucleatum were prepared in fluid thioglycollate medium (FTM) and incubated at 25 °C for 5-7 days. Two methods including conventional colony-forming unit (CFU) counting and direct microscopic count were used to evaluate cell viability. Average CFU number of each stain was determined by coating 100 μL microorganisms in exponential growth to tryptic soy agar (TSA) and repeated for at least 3 times. The blood counting chamber was used for direct microscopic count of microorganisms.

### NGSST assay

We developed standard operating procedures (SOPs) for NGS-based sterility test (NGSST). The NGSST assay workflow was performed as follows. Briefly, the whole genome DNA were amplified using the MGI Easy Single Cell Whole Genome Amplification Kit (Shenzhen, China) according to manufacturer’s instructions. Nuclear free water was used as negative control. The concentration and integrity of genomic DNA were evaluated by using Qubit and agarose gel electrophoresis. Then, the genomic DNA was purified using Agencourt AMPure XP (Beckman) and used to generate genomic DNA library. The whole genome sequencing, library construction and 100bp pair-end sequencing were conducted according to the protocol of DNBSEQ platform as described previously, and the acquired sequencing data of each sample was no less than 1Gb [13].

For sequence analysis, (1) Firstly, low-quality reads and adapters were filtered by Fastp software [14]. (2) The above filtered data (called clean reads) was aligned to the human reference genome hg38 by hisat2 [15]. (3) We then extracted the reads that mapped to the endogenous virus sequence of hg38, and (4) combined the reads mapped to the endogenous virus sequence with the reads not mapped to human reference genome hg38, and aligned them to the MetaPhlAn2 markers database [16]. (5) Finally, ! we analyzed the statistics of the above results, and calculated the number of reads of ä specific microorganisms per kilobase per million mapped reads (RPKM). The RPKM 1 ratio (RPKMr) for each microorganism was defined as:

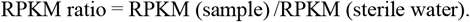

The minimum RPKM (sterile water) was set to 1.

### Establishment of detection parameters

The potential application of NGSST method was first verified in detecting microbial contamination under a given set of specific experimental conditions, and the differences in performance between the NGS-based method and the pharmacopoeia method were assessed.

To evaluate analytical performance of the NGSST, six model microorganisms at 0.1, 0.2, 0.5, 1, 5, 10, 50 and 100 CFU were prepared and tested by NGSST method. The RPKMr were calculated based on the alignment results from MetaPhlAn2 database, and limits of detection were evaluated. The ROC curve was drawn in R software (version 3.6.0) through pROC package to determine the 95% limits of detection (LOD). The criterion of RPMKr was established according to the ROC curve to report the detected bacteria or fungi. To evaluate the performance of NGSST in detecting early contamination during cell culture, 0.1, 0.2 and 0.5 CFU *E. coli* were incubated in DMEM-F12 cell culture medium supplied with 10% FBS for 14 days at 37°C, 5% CO2. The supernatant was then collected for sterility testing using both NGSST and BacT/ALERTR 3D methods.

### Statistical Analysis

To determine the optimal RPKM ratio threshold, we plotted the receiver operating characteristic (ROC) curves to determine the best RPKM ratio for the NGSST. The ROC curve was plotted with a confident level of 95%. The microorganisms that were added and detected by NGSST would be considered as truth values, while the microorganisms that were not added in the sample but detected would treat as false positive values. All statistical analysis was done in the R studio.

## Results

### Design of the NGSST development

Highly sensitive and rapid sterility testing plays an important role in biological product manufacturing such as T cells for adoptive transfer. We designed a novel sterility testing method based on next-generation sequencing. The schematic diagram of the NGSST shown in figure 1a includes whole genome amplification of samples, library construction and sequencing, alignment of reads, sterility testing report, and microorganism identification.

**Figure 1.**
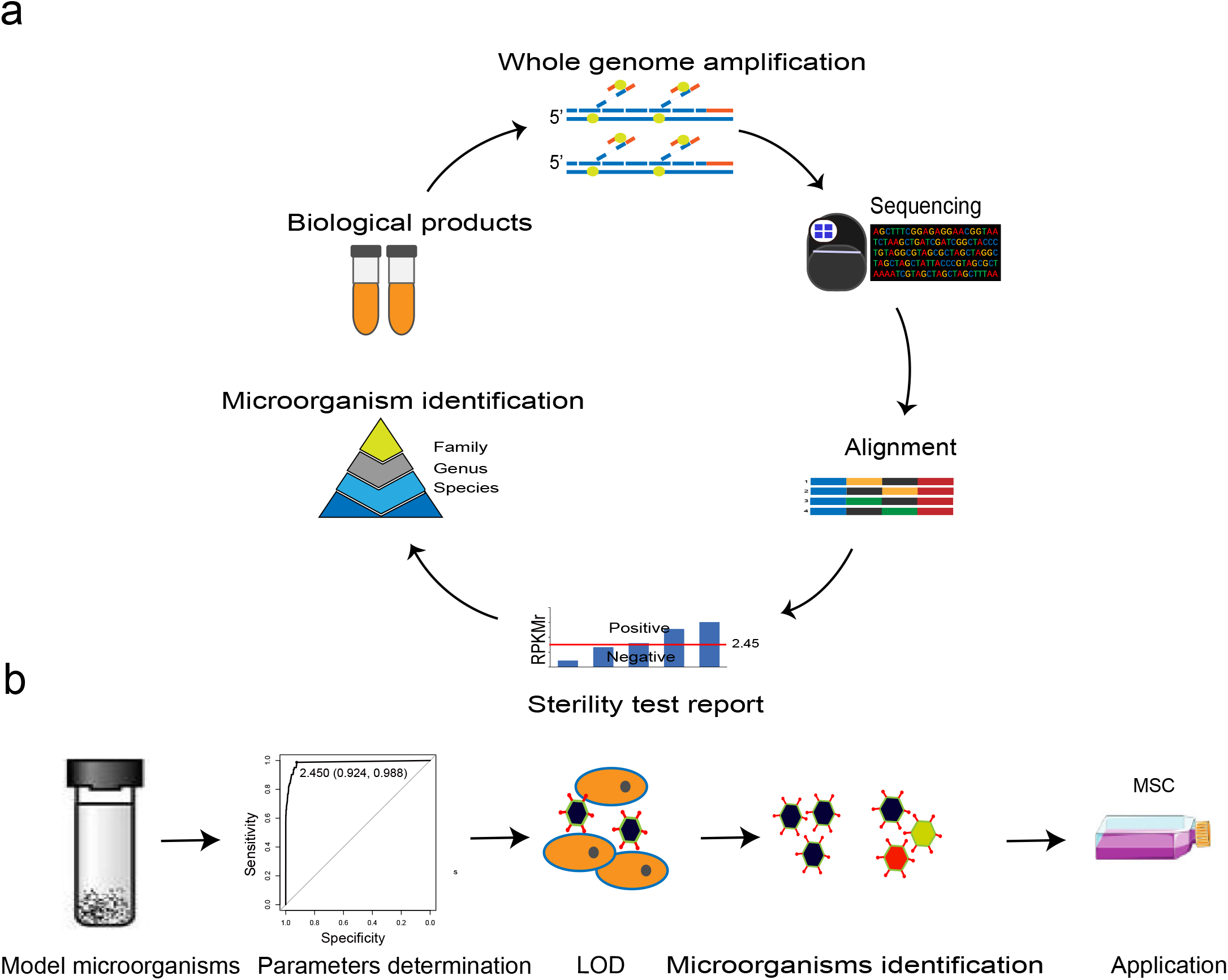
Design of the NGSST. **(a)** Schematic diagram of the NGSST on biological products. **(b)** Development and application of the NGSST. LOD, limit of detection.

To develop the NGSST, we followed four steps shown in Figure 1b. Firstly, we selected six model microorganisms (Table 1) including three bacteria strains with high GC content (*B. subtilis, E. coli*, and *P. aeruginosa*), two bacteria strains with low GC content (*C. sporogenes* and *S. aureus*), and fungi (*C. albicans*). We performed sequencing and reads mapping on marker genes of the six microorganisms. Secondly, we determined threshold to report positive results based on ROC (receiver operating characteristic) curve. Using the reads mapping data, we established limits of detection (the third step) and identified microorganisms (the fourth step). Finally, we finished the method development and applied it on biological samples. The sensitivity and accuracy of the NGSST have also been evaluated.

**Table 1.** Selected microorganisms in development and evaluation of the NGSST.

| Microorganisms | Strain ID | Anaerobic<br>/aerobic | Gram Stain | Incubation<br>temperature | GC-rich |
| --- | --- | --- | --- | --- | --- |
| Bacillus subtilis | CMCC(B)63501 | aerobic | Gram-positive | 30–35 | 44% |
| Candida albicans | CMCC(F)98001 | aerobic | Yeast | 20-25 | 39% |
| Clostridium sporogenes | CMCC(B)64941 | anaerobic | Gram-positive | 30–35 | 27% |
| Escherichia coli | ATCC 8739 | aerobic/an-<br>aerobic | Gram-negative | 30–35 | 51% |
| Pseudomonas<br>aeruginosa | CMCC(B)10104 | aerobic | Gram-negative | 30–35 | 54% |
| Staphylococcus aureus | CMCC(B)26003 | aerobic | Gram-positive | 30–35 | 33% |

### Development of the NGSST

We cultured the six microorganisms, counted colonies forming units (CFU) by their exponential growth on different agar plates, and diluted them into samples with microorganism concentrations at 0.1, 0.2, 0.5, 1, 10, 50, and 100 CFU. Since microorganisms in environment or solution may lead to false positive results, we prepared control samples with sterile water. With these samples as starting material, we conducted multiple displacement amplification (Table S1), libraries construction, and whole genome sequencing. The average sequencing data size was about 8 Gb (Table S2). We mapped reads onto microbial marker genes by using MetaPhAln2, calculated RPKM (reads per kilobase per million mapped reads), and computed RPKM ratio (RPKMr) for each microorganism as following:

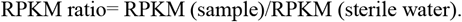

Most of sample’s RPKMrs were more than 100 (Figure 2a).

**Figure 2.**
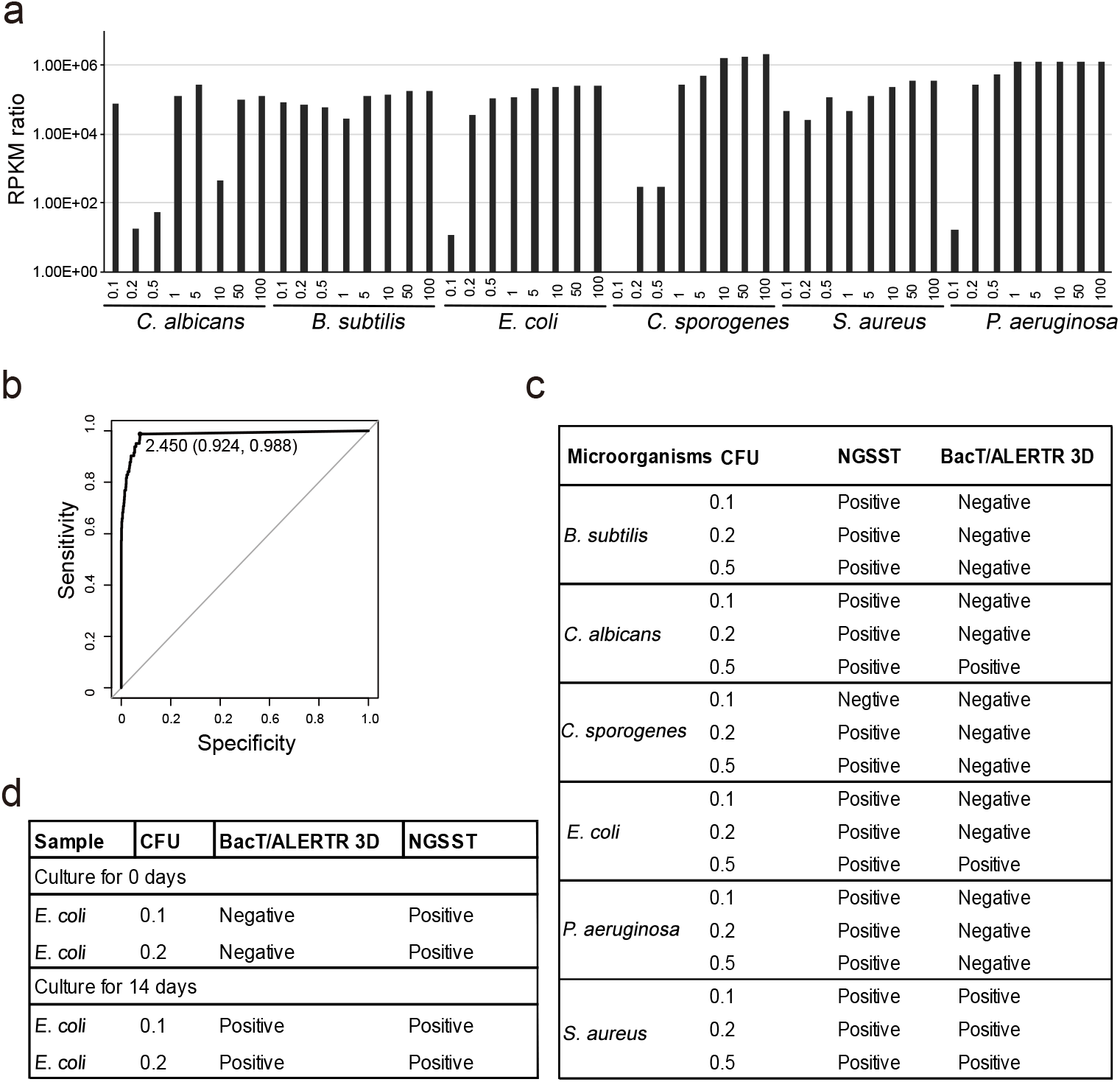
Establishment of the NGSST. **(a)** Whole genome sequencing and RPKM ratio of the six model microorganisms. Numbers on the X axis denote CFU (colonies forming units) of microorganisms through dilutions. **(b)** The determination of RPMKr threshold (RPMKr ≥2.45, AUC=0.983) based on ROC curve (specificity=0.924; sensitivity=0.988). **(c)** Higher sensitivity of the NGSST than that of BacT/ALERTR 3D. **(d)** Validations of the NGSST’s high sensitivity (0.1-0.2 CFU) by using culture compared with BacT/ALERTR 3D.

To determine RPKMr threshold, we established ROC curve and found that RPMKr ≥2.45 (AUC=0.983) would lead to high specificity and sensitivity (specifcity=0.924, sensitivity=0.988; Figure 2b). Using this threshold, we found that 47 out of all the 48 samples were detected postive of NGSST (Figure 2a). Interestingly, the NGSST showed high sensitivity that samples with microorganism concentration at 0.1CFU, 0.2CFU, and 0.5CFU could be detected by the NGSST (Figures 2a and 2c). Furthermore, we conducted sterility testing on these samples by using BacT/ALERTR 3D, and found that the NGSST (positive, 17 out of 18) displayed more accurate results than the BacT/ALERTR 3D (positive, 4 out of 18; Figure 2c). To validate this high sensitivity, we cultured the 0.1 and 0.2 CFU *E. coli* in medium for 14 days, and both sterility testing by NGSST and BacT/ALERTR 3D method displayed positive signal (Figure 2d).

### The application of NGSST on cell suspension

When the NGSST applied in mammalian cell suspension, plenty of nucleic acids released from mammalian cells appear to hinder the testing accuracy of the NGGST. To study the negative effect of human cells on the NGSST method, we prepared testing samples of MSC + *E. coli* by mixing human mesenchymal stem cells with 1, 5, 10, 50, and 100 CFU of *E. coli*. We conducted the NGSST which reported positive on all the MSC + *E. coli* samples (Figure 3a). However, the RPKMr values of MSC + *E. coli* samples were much smaller than that of *E. coli* samples.

**Figure 3.**
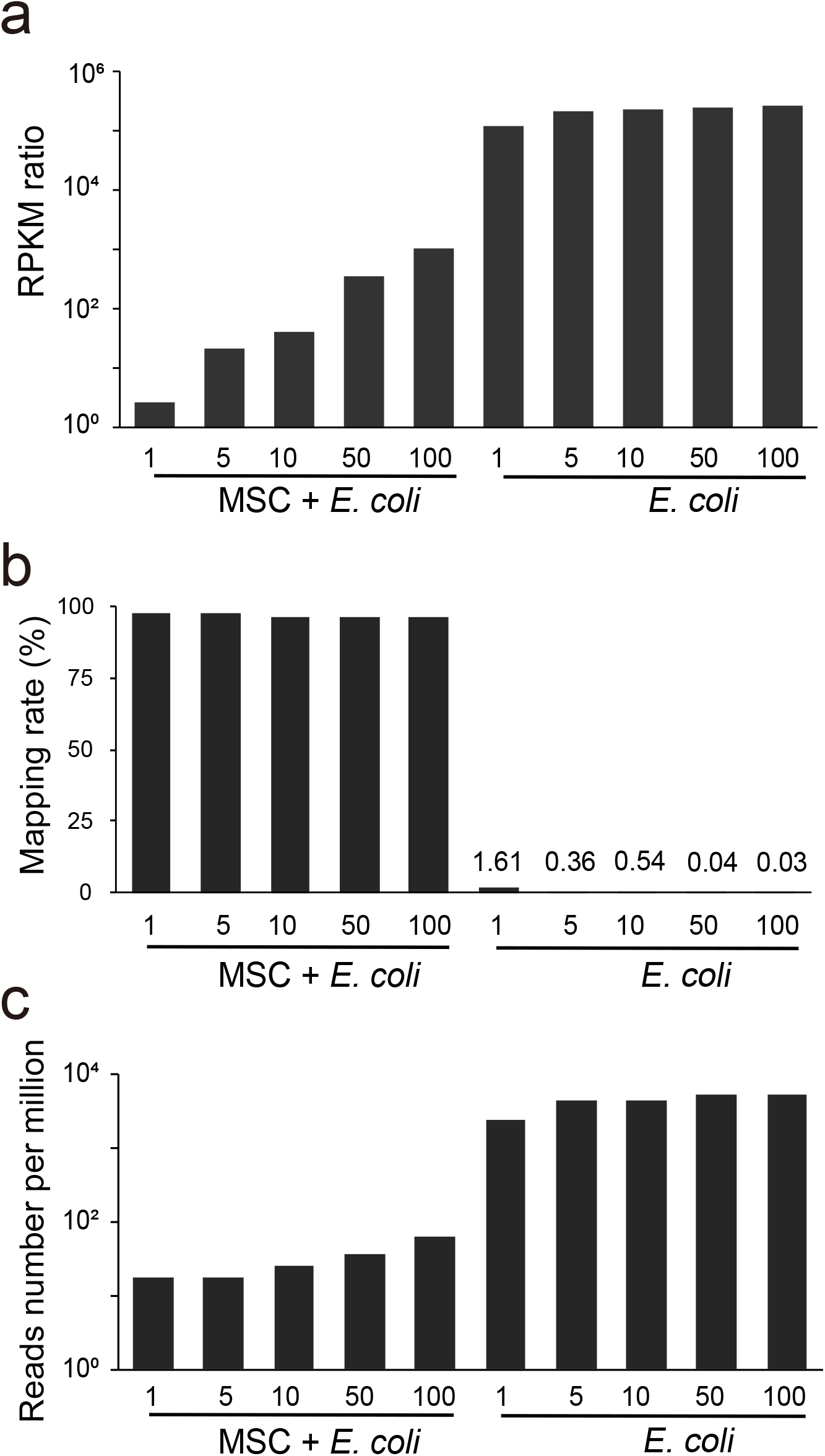
The NGSST LOD on the cell suspension mixed with *E. coli*. **(a)** The RPKMr values of the mixed suspension and *E. coli* samples. Numbers on the X axis denote CFU of microorganisms through dilutions. The human genome mapping rate **(b)** and marker reads per million mapped reads **(c)** in the mixed suspension and *E. coli* samples.

We analyzed sequencing reads by alignment to references of human genome or marker genes of MetaPhlAn2 database. The percentage of MSC + *E. coli* samples’ reads mapped to human genome reference (>95%) was around 50 folds than that of *E. coli* samples (<2%, Figure 3b), indicating that most of the sequencing reads were from human cells in MSC + *E.coli* samples. The values of reads mapped to MetaPhlAn2 database per million reads of MSC + *E. coli* samples (18 to 63) was much smaller than that of *E.coli* samples (>2000, Figure 3c). Overall, these results indicated that NGSST can be used in cell suspension with the LOD of 1 CFU. Compared with LOD of 0.1 CFU at *E. coli* group (Figure 2a), the sensitivity of sterility testing in MSC + *E. coli* samples was reduced due to most of the sequencing reads obtained from cultured cells.

### Identification of microorganisms by the NGSST

To assess the capability of the NGSST in identifying microorganism, we mixed two microorganisms (*E. coli* and *S. aureus*) and performed the NGSST. Results showed that *E. coli* or *S. aureus* were detected in all mixed samples of the two microorganisms (Figure 4a). Interestingly, *S. aureus* were identified with RPKMr of ~18 folds than the threshold in the mixture of 0.1 CFU *S. aureus* and 100 CFU *E. coli*. In addition, we prepared samples containing *E. coli*, *S. aureus, P. aeruginosa*, and human MSC. The *E. coli*, *S. aureus*, or *P. aeruginosa* were identified by the NGSST in all the samples even in the 1 CFU group (Figure 4b).

**Figure 4.**
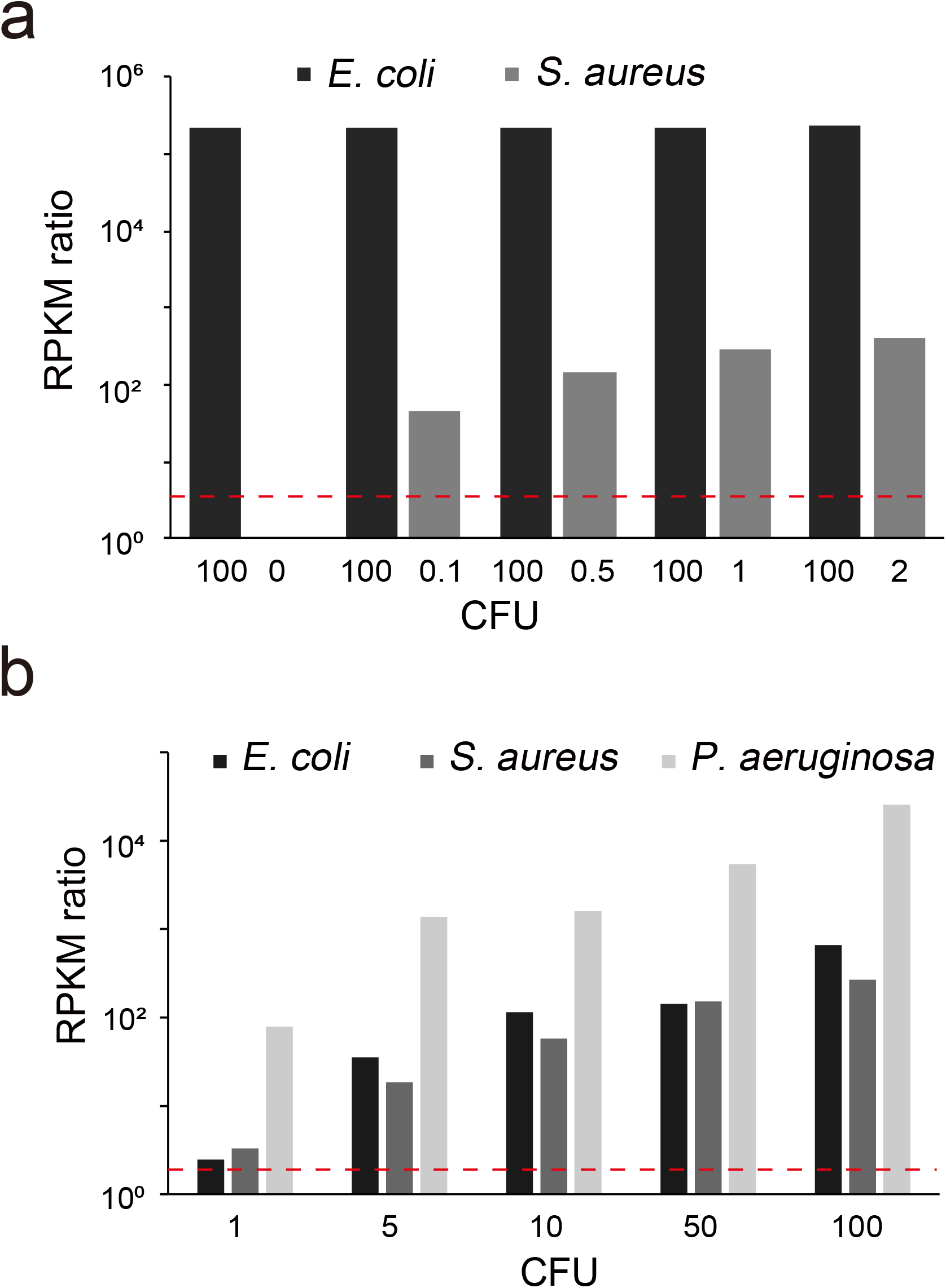
Identification of microorganisms by the NGSST. Detecting microorganism in two bacteria **(a)** or mixture of three bacteria and human MSC **(b)**. Red dotted line represents the NGSST threshold.

Furthermore, we applied the NGSST in five samples of cultured umbilical cord mesenchymal stem cells which were reported positive from BacT/ALERTR 3D testing. We identified several microorganisms in each of the five samples (Table 2). Interestingly, we also found that most of these identified bacteria strains are important components of vaginal microorganisms, which may be due to the contact between umbilical cord and vagina in vaginal birth [17]. Meanwhile, the total time required for NGSST method is roughly 24–48 hours, including whole genome amplication, sequencing, data analysis and interpretation, which is significantly shorter than current pharmacopoeia method (14 days).

**Table 2.** The microorganisms detected by NGSST on primary umbilical cord MSCs.

| Samples | Sterility testing by BacT/ALERTR 3D | Species | RPKMr |
| --- | --- | --- | --- |
| ucMSC-1 | Positive | *Bacteroides coprophilus* | 58.0 |
|  |  | *Erysipelotrichaceae bacterium* | 87.0 |
|  |  | *Lactobacillus iners* | 441.0 |
|  |  | *Ureaplasma parvum* | 1025.0 |
| ucMSC-2 | Positive | *Ureaplasma parvum* | 2584684.0 |
|  |  | *Ureaplasma urealyticum* | 18261.0 |
|  |  | *Xanthomonas perforans* | 22.9 |
| ucMSC-3 | Positive | *Acinetobacter sp NIPH* | 60.0 |
|  |  | *Bacteroides sp* | 49.0 |
|  |  | *Clostridium sp HGF2* | 118.0 |
|  |  | *Desulfovibrio sp* | 110.0 |
|  |  | *Erysipelotrichaceae bacterium* | 68.0 |
|  |  | *Lactobacillus iners* | 247.0 |
|  |  | *Parabacteroides sp* | 97.0 |
|  |  | *Ureaplasma parvum* | 472.0 |
|  |  | *Xanthomonas perforans* | 17.2 |
| ucMSC-4 | Positive | *Xanthomonas perforans* | 51.5 |
| ucMSC-5 | Positive | *Achromobacter xylosoxidans* | 13.3 |
|  |  | *Acinetobacter lwoffii* | 54.0 |
|  |  | *Afipia broomeae* | 169.0 |
|  |  | *Enterobacter cloacae* | 5560.0 |
|  |  | *Escherichia coli* | 209.3 |
|  |  | *Haemophilus aegyptius* | 178.0 |
|  |  | *Haemophilus haemolyticus* | 28.0 |
|  |  | *Haemophilus influenzae* | 36582.0 |
|  |  | *Klebsiella sp KTE92* | 3155.0 |
|  |  | *Propionibacterium acnes* | 462.0 |
|  |  | *Ralstonia solanacearum* | 26.0 |
|  |  | *Sphingomonas melonis* | 15.5 |
|  |  | *Staphylococcus warneri* | 128.1 |
|  |  | *Xanthomonas perforans* | 11.9 |

## Discussion

In this study, we developed and analytically validated the next-generation sequencing-based sterility test (NGSST) for biological products. Although the mNGS has been used for detection and characterization of pathogens in hospitalized patients [18], its application in sterility testing remained to be explored and optimized.

Currently, the available methods for rapid sterility testing include respiration based methods which use pharmacopoeia sterility testing and BacT/ALERT 3D system, and isolated DNA-based methods which involve nucleic acid amplification and whole-genome sequencing, as well as exogenous fluorescent substance related methods, such as solid phase cytometry and ATP bioluminescence [19–23]. Each method has its unique advantages and limitations with regard to sensitivity, time, and microbial identification.

NGSST method has many advantages in sterility test. The NGSST has a higher detection sensitivity (1 CFU), comparing to pharmacopoeial sterility test (100 CFU), ATP bioluminescence (1000 CFU), and solid phase cytometry (approximately 10 CFU). As to the time required for testing, pharmacopeia sterility test requires as long as 14 days, and the time required for NGSST method is about 48 hours, mainly involving whole genome amplification, DNA library construction, sequencing and data analysis. Furthermore, the NGSST is accurate for the identification of microorganisms. Although PCR amplification and sequencing of 16S ribosomal DNA also were used for microbial identification, the results are sometimes inaccurate due to the highly similar genomes between different microorganisms in 16S rDNA region. For example, although the DNA similarities between Mycobacterium chelonae and Mycobacterium abscessus is only 35%, the 16S r RNA has 99% homology. And other sterility testing methods lack the capability to identify microbial species.

The pipeline of NGSST method also overcomes certain challenges in application of whole genome sequencing in sterility testing. Most cell cultures or clinical samples may only contain small number of microorganisms, which makes it hard to directly extract sufficient DNA for sequencing. In 2014, Charles F. A. et al comparatively studied the performance of multiple displacement amplification (MDA), multiple annealing and looping based amplification cycles (MALBAC) and the PicoPLEX single cell WGA kit in single bacteria whole genome amplification, suggesting that a single bacterial cell with just 1 fg of genomic DNA is sufficient for WGA [24]. In NGSST method, we applied the MDA technology in amplification of bacteria and fungi genome, which effectively improves the detection sensitivity. Meanwhile, reproducibility threshold was established in the NGSST to correctly identify microorganisms from sequencing data and minimize false positive results. [25–27].

To avoid potential contaminations in the experimental process of WGA and DNA library construction, such as microorganisms from skin, environment and reagents, we utilized independent sterility testing room and ultraclean reagents with low level of DNA contamination, and as well set up conservative threshold values to minimize contamination [28].

Overall, we have explored the potential application of the NGSST in sterility testing of biological products, and demonstrated its high sensitivity, short detection period and capacity for microorganism identification. With continuously optimized cost of sequencing, the NGSST could be applied in in-process and product releasing stages of biological product manufacturing, as a supplementary to current pharmacopeia methods.

## Supporting information

Supplementary materia

## Declarations

### Ethics approval and consent to participate

This study was approved by the Institutional Review Board of BGI (No. BGI-IRB 19025-1-T1).

### Consent for publication

This Article is original, is not under consideration or has not been previous published elsewhere and its content has not been anticipated by any previous publication. All authors consent to publication of the article in Genome Research.

### Availability of data and material

The data that support the findings of this study have been deposited into CNGB Sequence Archive (CNSA) [29] of China National GeneBank DataBase (CNGBdb) [30] with accession number CNP0001476.

## Author’s contributions

X.Z., B.L. and J.Y., designed the study. J.Y., X.J., Q.M., performed the experiments. C.C. and B.L. were responsible for the data acquisition and data analysis. X.Z., B.L., and J.Y co-wrote the manuscript.

## Acknowledgements

We sincerely thank the support provided by China National GeneBank.

