## Supplementary materia for "Development of next-generation sequencing-based sterility test"

**Supplementary material**

**Figure S1**. Agarose gel electrophoresis of genomic DNA amplified by MDA.

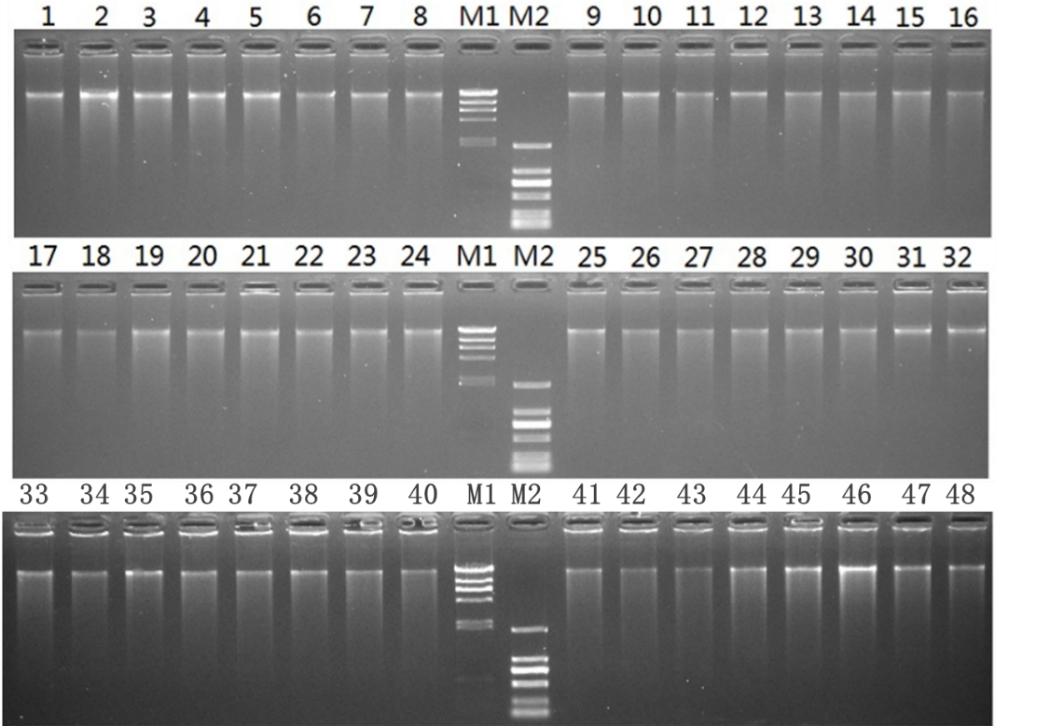

**Figure S1**. Agarose gel electrophoresis of genomic DNA amplified by MDA. 1-8: Genomics DNA from 0.1, 0.2, 0.5, 1, 5,10, 50, 100 CFU *P. aeruginosa*, 9-16: Genomics DNA from 0.1, 0.2, 0.5, 1, 5,10, 50, 100 CFU of *E. coli*, 17-24: Genomics DNA from 0.1, 0.2, 0.5, 1, 5,10, 50, 100 CFU *B. subtilis*. 25-32: Genomics DNA from 0.1, 0.2, 0.5, 1, 5,10, 50, 100 CFU *S. aureus*, 33-40: Genomics DNA from 0.1, 0.2, 0.5, 1, 5,10, 50, 100 CFU *C. albicans*, 41-48: Genomics DNA from 0.1, 0.2, 0.5, 1, 5,10, 50, 100 CFU *C. sporogenes*. M1: λ-Hind Ш digest ladder (Takara), M2: D2000 DNA ladder (Takara).

**Table S1** Quality Control of multiple displacement amplification.

| **Microorganisms** | **CFU** | **DNA concentration (ng/µL)** | **Volume (µL)** | **Total mass (µg)** |
| --- | --- | --- | --- | --- |
| *B. subtilis* | 0.1 | 194 | 42 | 8.1 |
|  | 0.2 | 185 | 42 | 7.8 |
|  | 0.5 | 172 | 42 | 7.2 |
|  | 1 | 286 | 23 | 6.6 |
|  | 5 | 212 | 23 | 4.9 |
|  | 10 | 264 | 23 | 6.1 |
|  | 50 | 356 | 23 | 8.2 |
|  | 100 | 366 | 23 | 8.4 |
| *C. albicans* | 0.1 | 174 | 42 | 7.3 |
|  | 0.2 | 194 | 42 | 8.1 |
|  | 0.5 | 189 | 42 | 7.9 |
|  | 1 | 137 | 42 | 5.8 |
|  | 5 | 120 | 42 | 5.0 |
|  | 10 | 228 | 23 | 5.2 |
|  | 50 | 193 | 23 | 4.4 |
|  | 100 | 148 | 23 | 3.4 |
| *C. sporogenes* | 0.1 | 208 | 42 | 8.7 |
|  | 0.2 | 194 | 42 | 8.1 |
|  | 0.5 | 183 | 42 | 7.7 |
|  | 1 | 242 | 23 | 5.6 |
|  | 5 | 300 | 23 | 6.9 |
|  | 10 | 196 | 23 | 4.5 |
|  | 50 | 147 | 23 | 3.4 |
|  | 100 | 137 | 23 | 3.2 |
| *E. coli* | 0.1 | 197 | 42 | 8.3 |
|  | 0.2 | 212 | 42 | 8.9 |
|  | 0.5 | 260 | 42 | 10.9 |
|  | 1 | 204 | 23 | 4.7 |
|  | 5 | 360 | 15 | 5.4 |
|  | 10 | 416 | 23 | 9.6 |
|  | 50 | 450 | 23 | 10.4 |
|  | 100 | 608 | 23 | 14.0 |
| *P. aeruginosa* | 0.1 | 208 | 42 | 8.7 |
|  | 0.2 | 238 | 42 | 10.0 |
|  | 0.5 | 296 | 42 | 12.4 |
|  | 1 | 564 | 23 | 13.0 |
|  | 5 | 774 | 23 | 17.8 |
|  | 10 | 592 | 23 | 13.6 |
|  | 50 | 584 | 23 | 13.4 |
|  | 100 | 702 | 23 | 16.1 |
| *S. aureus* | 0.1 | 200 | 42 | 8.4 |
|  | 0.2 | 186 | 42 | 7.8 |
|  | 0.5 | 172 | 42 | 7.2 |
|  | 1 | 170 | 23 | 3.9 |
|  | 5 | 184 | 23 | 4.2 |
|  | 10 | 224 | 23 | 5.2 |
|  | 50 | 210 | 23 | 4.8 |
|  | 100 | 234 | 23 | 5.4 |

**Table S2** Sequencing data summary.

| **Sample** | **CFU** | **Raw base (G)** | **Clean base (G)** | **Clean rate** | **Raw Q20** | **Clean Q20** | **Raw Q30** | **Clean Q30** |
| --- | --- | --- | --- | --- | --- | --- | --- | --- |
| Control (water) | 0 | 13.36 | 12.87 | 0.963 | 0.97 | 0.98 | 0.9 | 0.91 |
| *C. albicans* | 0.1 | 9.82 | 9.38 | 0.955 | 0.97 | 0.97 | 0.89 | 0.9 |
|  | 0.2 | 8.93 | 8.55 | 0.957 | 0.97 | 0.97 | 0.89 | 0.9 |
|  | 0.5 | 9.39 | 8.95 | 0.953 | 0.97 | 0.97 | 0.88 | 0.89 |
|  | 1 | 10.46 | 10.04 | 0.96 | 0.97 | 0.98 | 0.9 | 0.91 |
|  | 5 | 10.06 | 9.67 | 0.961 | 0.97 | 0.98 | 0.9 | 0.91 |
|  | 10 | 8.06 | 7.89 | 0.979 | 0.98 | 0.99 | 0.93 | 0.93 |
|  | 50 | 8.7 | 8.57 | 0.985 | 0.99 | 0.99 | 0.93 | 0.93 |
|  | 100 | 6.09 | 5.8 | 0.952 | 0.97 | 0.98 | 0.89 | 0.9 |
| *E. coli* | 0.1 | 10.35 | 9.96 | 0.962 | 0.97 | 0.98 | 0.9 | 0.9 |
|  | 0.2 | 15.04 | 14.37 | 0.955 | 0.97 | 0.97 | 0.89 | 0.9 |
|  | 0.5 | 12.4 | 11.9 | 0.96 | 0.97 | 0.98 | 0.89 | 0.9 |
|  | 1 | 7.31 | 7.19 | 0.984 | 0.99 | 0.99 | 0.93 | 0.93 |
|  | 5 | 7.06 | 6.92 | 0.98 | 0.98 | 0.99 | 0.92 | 0.93 |
|  | 10 | 6.47 | 6.36 | 0.983 | 0.99 | 0.99 | 0.93 | 0.94 |
|  | 50 | 7.44 | 7.3 | 0.981 | 0.98 | 0.98 | 0.92 | 0.92 |
|  | 100 | 6.08 | 5.99 | 0.985 | 0.99 | 0.99 | 0.93 | 0.93 |
| *S. aureus* | 0.1 | 10.43 | 10.02 | 0.961 | 0.97 | 0.98 | 0.9 | 0.9 |
|  | 0.2 | 11.56 | 11.09 | 0.959 | 0.97 | 0.98 | 0.89 | 0.9 |
|  | 0.5 | 12.6 | 12.13 | 0.963 | 0.97 | 0.98 | 0.9 | 0.91 |
|  | 1 | 7.85 | 7.72 | 0.983 | 0.98 | 0.99 | 0.93 | 0.93 |
|  | 5 | 7.63 | 7.51 | 0.984 | 0.99 | 0.99 | 0.93 | 0.93 |
|  | 10 | 9.65 | 9.5 | 0.984 | 0.99 | 0.99 | 0.93 | 0.93 |
|  | 50 | 6.55 | 6.48 | 0.989 | 0.99 | 0.99 | 0.94 | 0.94 |
|  | 100 | 7.6 | 7.5 | 0.987 | 0.99 | 0.99 | 0.93 | 0.93 |
| *B. subtilis* | 0.1 | 11.06 | 10.56 | 0.955 | 0.97 | 0.97 | 0.88 | 0.9 |
|  | 0.2 | 11.76 | 11.29 | 0.96 | 0.97 | 0.98 | 0.9 | 0.91 |
|  | 0.5 | 9.37 | 8.98 | 0.958 | 0.97 | 0.97 | 0.89 | 0.9 |
|  | 1 | 5.56 | 5.45 | 0.98 | 0.98 | 0.99 | 0.92 | 0.93 |
|  | 5 | 6.18 | 6.06 | 0.981 | 0.98 | 0.99 | 0.92 | 0.92 |
|  | 10 | 8.19 | 8.05 | 0.983 | 0.98 | 0.99 | 0.92 | 0.93 |
|  | 50 | 7.39 | 7.28 | 0.985 | 0.99 | 0.99 | 0.93 | 0.94 |
|  | 100 | 8.22 | 8.08 | 0.983 | 0.98 | 0.99 | 0.92 | 0.93 |
| *C. sporogenes* | 0.1 | 11.27 | 10.8 | 0.958 | 0.97 | 0.98 | 0.89 | 0.9 |
|  | 0.2 | 12.12 | 11.61 | 0.958 | 0.97 | 0.98 | 0.89 | 0.9 |
|  | 0.5 | 8.81 | 8.37 | 0.95 | 0.97 | 0.97 | 0.88 | 0.89 |
|  | 1 | 6.62 | 6.28 | 0.949 | 0.97 | 0.97 | 0.89 | 0.9 |
|  | 5 | 6.95 | 6.63 | 0.954 | 0.97 | 0.98 | 0.89 | 0.9 |
|  | 10 | 2.08 | 1.99 | 0.957 | 0.97 | 0.98 | 0.89 | 0.9 |
|  | 50 | 3.63 | 3.47 | 0.956 | 0.97 | 0.98 | 0.9 | 0.91 |
|  | 100 | 5.03 | 4.84 | 0.962 | 0.98 | 0.98 | 0.91 | 0.92 |
| *P. aeruginosa* | 0.1 | 11.28 | 10.81 | 0.958 | 0.97 | 0.98 | 0.9 | 0.91 |
|  | 0.2 | 12.1 | 11.55 | 0.955 | 0.97 | 0.97 | 0.89 | 0.9 |
|  | 0.5 | 10.54 | 9.83 | 0.933 | 0.96 | 0.97 | 0.87 | 0.88 |
|  | 1 | 6.61 | 6.4 | 0.968 | 0.98 | 0.98 | 0.9 | 0.91 |
|  | 5 | 3.85 | 3.7 | 0.961 | 0.97 | 0.98 | 0.89 | 0.89 |
|  | 10 | 6.26 | 6 | 0.958 | 0.97 | 0.98 | 0.88 | 0.89 |
|  | 50 | 6.03 | 5.81 | 0.964 | 0.97 | 0.98 | 0.89 | 0.9 |
|  | 100 | 6.15 | 5.93 | 0.964 | 0.97 | 0.98 | 0.89 | 0.9 |
| *E. coli* + *S. aureus* | 0 | 4.05 | 3.93 | 0.97 | 0.98 | 0.98 | 0.91 | 0.92 |
|  | 0.1 | 5.49 | 5.34 | 0.973 | 0.98 | 0.98 | 0.92 | 0.92 |
|  | 0.5 | 4.08 | 3.95 | 0.968 | 0.98 | 0.98 | 0.91 | 0.92 |
|  | 1 | 4.52 | 4.38 | 0.969 | 0.98 | 0.98 | 0.91 | 0.92 |
|  | 2 | 3.82 | 3.7 | 0.969 | 0.98 | 0.98 | 0.91 | 0.91 |
| MSC +*E. coli* + *S. aureus* + *P. aeruginosa* | 1 | 31.74 | 30.96 | 0.975 | 0.98 | 0.99 | 0.93 | 0.93 |
|  | 5 | 14.08 | 13.74 | 0.976 | 0.98 | 0.98 | 0.91 | 0.92 |
|  | 10 | 19.71 | 19.33 | 0.981 | 0.98 | 0.99 | 0.93 | 0.93 |
|  | 50 | 13.02 | 12.74 | 0.978 | 0.98 | 0.99 | 0.92 | 0.93 |
|  | 100 | 13.53 | 13.22 | 0.977 | 0.98 | 0.98 | 0.92 | 0.92 |
| MSC + *E. coli* | 1 | 16.83 | 16.44 | 0.977 | 0.98 | 0.98 | 0.92 | 0.92 |
|  | 5 | 18.43 | 18.07 | 0.98 | 0.98 | 0.99 | 0.92 | 0.93 |
|  | 10 | 18.62 | 17.87 | 0.96 | 0.98 | 0.99 | 0.93 | 0.93 |
|  | 50 | 14.99 | 14.66 | 0.978 | 0.99 | 0.99 | 0.93 | 0.94 |
|  | 100 | 20.07 | 19.64 | 0.979 | 0.98 | 0.99 | 0.92 | 0.93 |
| ucMSC-1 | NA | 9.37 | 9.11 | 0.972 | 0.98 | 0.98 | 0.9 | 0.91 |
| ucMSC-2 | NA | 10.6 | 10.38 | 0.979 | 0.98 | 0.99 | 0.93 | 0.93 |
| ucMSC-3 | NA | 10.44 | 10.18 | 0.975 | 0.98 | 0.98 | 0.92 | 0.92 |
| ucMSC-4 | NA | 8.54 | 8.17 | 0.957 | 0.97 | 0.98 | 0.89 | 0.9 |
| ucMSC-5 | NA | 17.17 | 16.78 | 0.977 | 0.98 | 0.98 | 0.92 | 0.92 |

Note: MSC, mesenchymal stem cells; ucMSC, umbilical cord mesenchymal stem cells.
